# Profiling age and body fluid DNA methylation markers using nanopore adaptive sampling

**DOI:** 10.1101/2023.12.10.571037

**Authors:** Zaka Wing-Sze Yuen, Somasundhari Shanmuganandam, Maurice Stanley, Simon Jiang, Nadine Hein, Runa Daniel, Dennis McNevin, Cameron Jack, Eduardo Eyras

## Abstract

DNA methylation plays essential roles in regulating physiological processes, from tissue and organ development to gene expression and aging processes and has emerged as a widely used biomarker for the identification of body fluids and age prediction. Currently, methylation markers are targeted independently at specific CpG sites, as part of a multiplexed assay, rather than through a unified assay. Methylation detection is also dependent on divergent methodologies, ranging from enzyme digestion and affinity enrichment to bisulfite treatment, alongside various technologies for high-throughput profiling, including microarray and sequencing. In this pilot study, we test the simultaneous identification of age-associated and body fluid-specific methylation markers using a single technology, nanopore adaptive sampling. This innovative approach enables the profiling of multiple CpG marker sites across entire gene regions from a single sample without the need for specialized DNA preparation or additional biochemical treatments. Our study demonstrates that adaptive sampling achieves sufficient coverage in regions of interest to accurately determine the methylation status, shows a robust consistency with whole-genome bisulfite sequencing data, and corroborates known CpG markers of age and body fluids. Our work also resulted in the identification of new sites strongly correlated with age, suggesting new possible age methylation markers. This study lays the groundwork for the systematic development of nanopore-based methodologies in both age prediction and body fluid identification, highlighting the feasibility and potential of nanopore adaptive sampling while acknowledging the need for further validation and expansion in future research.

## Introduction

DNA methylation is a significant epigenetic modification in genomes, involving the addition of methyl groups to specific nucleotides. In humans, the prevailing form of DNA methylation is 5-methylcytosine (5mC), primarily found at CpG dinucleotides, which are often concentrated in extensive clusters known as CpG islands (CGIs). The addition and removal of methyl groups are mediated by enzymes such as DNA methyltransferases (DNMTs) and Ten-eleven translocation (TET) enzymes, respectively, allowing the reversibility of DNA methylation. This dynamic nature is essential for maintaining genome stability, regulating gene expression, and responding to environmental changes throughout mammalian development and throughout an organism’s lifespan (Greenberg & Bourc’his, 2019; Jones et al., 2015; Jones, 2012; Kader & Ghai, 2015; Raiber et al., 2017). DNA methylation undergoes significant alterations during development, starting from the germline and early embryo, where extensive demethylation occurs to reset the epigenetic state, followed by the reconstitution of methylation around the time of implantation (Greenberg & Bourc’his, 2019). For instance, DNA methylation is fundamental in the inactivation of one X chromosome in females, a vital process for ensuring comparable levels of X-linked gene expression in both sexes (Greenberg & Bourc’his, 2019). Moreover, DNA methylation is influenced by environmental factors and varies among different tissues, cell types, and disease states (Greenberg & Bourc’his, 2019). For example, irregularities in DNA methylation patterns, such as the hypermethylation or hypomethylation of genes, can lead to inappropriate gene silencing or activation, respectively, thereby contributing to the onset and progression of diseases like cancer.

Since DNA methylation varies across cell types and tissues, and throughout an organism’s lifetime, it has been identified as one of the most promising biomarkers for age prediction and body fluid identification in forensic applications. Indeed, DNA methylation detection can be applied as part of forensic DNA phenotyping (FDP) to provide intelligence by inferring the age of a person to assist in an investigation (Dias et al., 2020; Freire-Aradas et al., 2016; Freire-Aradas et al., 2020; Jones et al., 2015; Lee et al., 2022; Montesanto et al., 2020; Vidaki et al., 2017; Woźniak et al., 2021). Numerous age prediction models (APMs) based on DNA methylation have been developed to infer the biological age of individuals, serving as a proxy for chronological age. Some recent models are based on hundreds of CpG sites using a genome-wide microarray (Bocklandt et al., 2011; Hannum et al., 2013; Horvath, 2013; Levine et al., 2018). A recent development introduced a universal pan-mammalian clock using methylation arrays, spanning 59 tissue types across 185 mammalian species, and exhibiting a high correlation with age (r > 0.96) (Lu et al., 2023). However, microarrays require more DNA than is typically recovered from biological evidence which restricts their practical application in a forensic context.

Given these limitations, tissue-specific APMs have been commonly used which focus on a small set of age-associated CpG sites using a broad spectrum of targeted approaches. Widely used technologies to detect methylation in these APMs include Sanger sequencing (Dias et al., 2020), pyrosequencing (Bekaert et al., 2015; Feng et al., 2018; Fleckhaus & Schneider, 2020; Li et al., 2020; Park et al., 2016; Thong et al., 2017; Weidner et al., 2014; Zbieć-Piekarska, Spólnicka, Kupiec, Makowska, et al., 2015; Zbieć-Piekarska, Spólnicka, Kupiec, Parys-Proszek, et al., 2015), SNaPshot (Hong et al., 2017; Jung et al., 2019; Lee et al., 2022), massively parallel sequencing (MPS) (Aliferi et al., 2018; Naue et al., 2017; Naue et al., 2018; Vidaki et al., 2017; Vidaki & Kayser, 2018; Woźniak et al., 2021) and EpiTYPER mass spectrometry (Freire-Aradas et al., 2018; Freire-Aradas et al., 2016). Two key characteristics common to all these forensic APMs are that they only incorporate a few of the most statistically reliable CpG sites into the model and that they require at least 20 ng DNA input to ensure robust methylation quantification. While less than required by microarrays, this DNA input requirement is still higher than the amount of DNA typically recovered from crime scenes or available after established forensic STR-based methodologies have been applied.

Various techniques have also been established for the identification of body fluids, encompassing the analysis of RNA expression specific to certain cell types (Fleming & Harbison, 2010a; Haas et al., 2009; Hanson et al., 2009; Ingold et al., 2018; Juusola & Ballantyne, 2003, 2007; Lindenbergh et al., 2012; Salzmann et al., 2021; Zubakov et al., 2010), the detection of tissue-specific microRNA profiles (Coenen-Stass et al., 2018; Silva et al., 2015; Weber et al., 2010), the detection of DNA methylation unique to particular tissues (Alghanim et al., 2020; Frumkin et al., 2011; Gauthier et al., 2019; Jung et al., 2016; Lee et al., 2015; Lee et al., 2012; Lee et al., 2022; Madi et al., 2012), protein profiling (de Beijer et al., 2018; Legg et al., 2014, 2017; Van Steendam et al., 2013; Yang et al., 2013), and the identification of microbiomes residing within tissues (Dobay et al., 2019; Fleming & Harbison, 2010b; Hanssen et al., 2017; Salzmann et al., 2019).

Another common characteristic of the current assays for age prediction and body fluid identification is their reliance on bisulfite conversion for the profiling of 5mC. However, the disadvantages of this process include substantial DNA degradation, incomplete conversion, reduced sequence variability, and the necessary coupling with lengthy protocols (Freire-Aradas et al., 2020; Grunau et al., 2001; Yong et al., 2016). Furthermore, the amount of bisulfite-converted DNA required for polymerase chain reaction (PCR) and systematic errors associated with the analytical processes used in capillary electrophoresis (CE) analysis from the SNaPshot assays can influence DNA methylation quantification (Hong & Shin, 2021; Lee et al., 2022). Furthermore, the detection of methylation markers requires the inclusion of each locus of interest in a multiplexed assay. While numerous CpG sites have been recognized and used for age prediction and body fluid identification, detecting methylation markers for each purpose are commonly conducted separately in each experiment, thereby increasing the total number of assays required.

To address these challenges, this study utilised nanopore sequencing which allows the direct sequencing of native DNA molecules carrying base modifications, notably 5mC, without requiring any additional sample processing beyond standard library preparation. Given that the average nanopore reads lengths are now >20,000 bp (20 kb), this technology is ideal for sequencing both small targets, including STRs and SNPs and CpG sites, as well as significant portions of longer haplotypes at a fraction of the cost of MPS-based approaches (Hall et al., 2020; Plesivkova et al., 2019; Wang et al., 2021).This approach avoids depletion of DNA retrieved from biological evidence, a consequence of multiple subsequent analysis undertaken after STR profiling to assist in identify the DNA donor. To efficiently test for multiple methylation markers with the same technology, we used the adaptive sampling strategy, also known as ReadUntil. This approach combines real-time basecalling with rapid mapping to selectively identify and discard off-target reads during the sequencing process. A fragment that enters a pore will produce sequence data in real time and is assessed for proximity to a target by rapid alignment to a reference gnome. If the target is sufficiently close, the remainder of the fragment is sequenced. If it is not, the fragment is ejected from the pore and another fragment is allowed access. In this way, unwanted sequence data is minimised and coverage can be focussed on regions of interest (ROIs) (Martin et al., 2022).

By programming these devices to recognize specific DNA molecules, computational enrichment of ROIs becomes possible, eliminating the need for specialized reagents. We used nanopore adaptive sampling to corroborate existing forensic age-associated markers and identify potential new novel markers by investigating all CpG sites within the target genes. This study demonstrates the applicability of this approach, and streamlined sample processing, to the detection and analysis of age methylation markers, establishing the basis for the broader applicability of nanopore adaptive sampling in forensic casework and age-related studies.

## Methods

### DNA samples

Three different types of gDNA samples were used in the adaptive sampling runs. Firstly, the GIAB reference DNA sample HG002 was employed to evaluate the performance of our custom-designed target panel.

Secondly, we conducted two adaptive sampling runs using methylation control samples from EpigenDX (https://www.epigendx.com/d/products/methylation-controls). These methylation controls were derived from human whole blood and enzymatically treated, resulting in the high methylated control with over 85% methylation and the low methylated control with less than 5% methylation, according to the manufacturer (EpigenDX, 2018).

Lastly, we tested adaptive sampling on human blood samples obtained with approval from the ethics committee from the Australian National University (ANU) under ethics protocol no. ETH.1.16.01/ETH.01.15.015. Blood samples were collected at the Centre for Personalised Immunology (ANU) from ten participants aged 25 to 76 years. DNA extraction from the blood samples was performed using the Monarch HMW DNA Extraction Kit for Cells & Blood (NEB, T3050) following the manufacturer’s protocol (New England Biolabs, 2023). The age and biological gender of the ten donor samples are listed in Supp. Table 1.

### Quality control

The concentration of the gDNA samples was measured with the NanoDrop based on A260 while the purity was assessed using A260:A280 and A260:A230 ratios. Genomic DNA concentration was also determined using the Qubit dsDNA Broad Range Assay Kit (Thermo Fisher Scientific). The Agilent Femto Pulse system was also used to assess the fragment size distribution of the samples, and thus determine suitability for shearing and size selection in the next step.

### Shearing and size selection

Based on the initial QC results, which indicated that that DNA samples were excessively condensed and clogged, the samples underwent a needle-shearing process to make them easier to pipette. The samples were needle-sheared 5 to 15 times with 26 and/or 29 gauge needles, followed by confirmation on the Agilent Femto Pulse system. The samples were then electrophoretically size-selected using the BluePippin system (Sage science, USA) with 0.75% Agarose gel cassettes and marker S1, enriching for fragments that were at least 20kb long. Samples were then recovered and further size-selected with AMPure XP beads and quantified using Qubit with dsDNA Broad Range reagents to assess suitability for subsequent library preparation. The fragment sizes of the enriched and purified samples, ranging from 20 to 50 kb, were confirmed by Femto Pulse.

### Library preparation and sequencing with adaptive sampling

Standard Nanopore sequencing libraries were prepared from genomic DNA methylation controls purchased from EpigenDx, using SQK-LSK110 gDNA by ligation sequencing kit (ONT) following the manufacturer’s instructions. Details regarding the input DNA amounts and the final DNA amounts and concentrations obtained post-library preparation, are listed in Supp. Table 1. Each resulting library was then loaded into a MinION flow cell (FLO-MIN106D, R9.4.1 pore) and sequenced on the MinION sequencer which was connected via a Type-A USB 3.0 port on a computer workstation with the NVIDIA GeForce RTX 3070 GPU card. MinKNOW version v20.10.3 and Guppy version v4.2.2 were utilised (Oxford Nanopore Technologies, 2023b).

Using the MinKNOW software, an adaptive sampling run (set to enrich) was performed with sequencing parameters set to use the ‘fast’ basecalling model of Guppy (Oxford Nanopore Technologies, 2023a). Target regions were defined as a multi-fasta file including forensically relevant STR and SNP loci, 10 genes commonly employed in forensic age prediction models, 307 aging-related human genes from the Ageing Gene Database (Tacutu et al., 2018), 393 orthologous genes displaying conserved time-dependent behavior across mammals (Wang et al., 2020), 31 genes associated with body fluid identification, and three copies of a tandem repeat of the rDNA unit (KY962518.1) containing two units in three possible orientations with respect to the direction of transcription: 1) the same orientation, 2) convergent or 3) divergent. Additional information indicating the targeted regions can be found in Supp. Table 2.

The inclusion of 393 orthologous genes in our study is mainly driven by their shared epigenetic signature, which mirrors the developmental stages in dogs, humans, and mice. This characteristic makes them suitable for creating a universal epigenetic clock of aging across species. Consequently, we have also added these orthologues to our target panel to explore the possibility of identifying additional genes linked to aging in humans. In addition to the orthologous genes, our study also aims to investigate the accuracy of CpG methylation detection using nanopore adaptive sampling. To this end, we targeted regions that include three copies of a tandem repeat of the rDNA unit, each comprising two units in varied orientations. This approach enables us to utilize the adaptive sampling technology not only to enrich and sequence individual rDNA copies but also to assess their methylation status, employing the same control samples used for methylation verification.

Most samples were completed within 48 hours, with nuclease flushing of the flow cell to increase throughput and additional loading to the washed flow cell. Based on sequencing output, samples that produced less than two million reads required additional library preparation. These were run for another ∼36 hours, at which point the nanopores in the flow cells were too damaged or disabled to generate further data.

### Data processing and methylation analysis

Nanopore reads in FAST5 format were analysed with Bonito (v0.7.2) using a Remora (v2.1.3) model (dna_r9.4.1_e8_sup_5mC) to perform modified basecalling and alignment to the modified GRCh38/hg38 human genome assembly containing three additional tandem copies of the rDNA canonical unit (KY962518.1). ModBAM files with MM and ML tags generated by Bonito were then converted into bedMethyl files using Modkit (v0.1.13) pileup (https://github.com/nanoporetech/modkit) with the traditional preset. The bedMethyl file aggregated data across strands, providing information on counts and the proportion of calls classified as modified at CpG sites. The following columns from the resulting bedMethyl files were extracted: chromosome, start position, end position, coverage and methylation fraction, to reduce the file size and prepare for downstream analyses in R (v4.3.1). These data were then aggregated from ten blood donor samples.

A simple linear regression analysis was conducted on the combined dataset of all CpG sites identified from the sequencing data. This analysis examined the relationship between methylation level of each CpG sites and chronological age, yielding a correlation coefficient (*r*), an associated *p*-value, and the RMSE for each CpG site. To improve precision we excluded CpG sites located on sex chromosomes to negate potential confounding effects due to possible differences in gene methylation that can be sex dependent (Gershoni & Pietrokovski, 2017).

A meta-analysis using the unweighted Stouffer’s method (Lu et al., 2023) was performed to synthesize collective statistical evidence across all CpG sites, which combined p-values generated from the simple linear regression to produce a meta Z-score and a meta *p*-value for each CpG site. By establishing a stringent threshold for the meta *p*-value at 0.00001, we identified CpG sites that demonstrated highly significant correlations with age.

Using the same aggregated datasets derived from ten different blood samples, nine body-specific CpG sites were selectively filtered, with their details provided in Supp. Table 3. These identified markers were employed for the purpose of body fluid identification (Lee et al., 2022). The methylation levels for these specific markers were then plotted.

## Results

### Adaptive sampling efficiently enriches genomic regions of interest

The effectiveness of adaptive sampling using a combination of regions, including 10 genes commonly employed in forensic age prediction models, 307 aging-related human genes (Tacutu et al., 2018), 393 genes displaying conserved time-dependent behavior across mammals (Wang et al., 2020), 31 genes associated with body fluid identification (Lee et al., 2022), and a tandem repeat of the rDNA unit, whose methylation has also been linked to aging in blood samples (Wang & Lemos, 2019) was evaluated. Our adaptive sampling approach covered the entire locus for each gene, including 20 kb of flanking sequencing in either direction. Overall, the panel encompassed approximately 113 Mb, targeting about 3.65% of the human reference genome (hg38) (Supp. Table 2).

To assess the target enrichment performance, adaptive sampling with our target panel was applied on the genome in a bottle (GIAB) reference sample HG002. Accepted reads displayed a bimodal distribution with two peaks, with one matching the desired target size of approximately 40 kb, and the other indicating the presence of short reads that could either be short on-target reads or false classifications (Figure 1a). In contrast, rejected reads exhibited a sharp and narrow peak around 500 bases, which is the length selected to eject the molecules from the pore when they are identified as not being from any target region. Additionally, there were reads with size ranging from 100 to about 2,500 bases, which the software could not classify, therefore, these fragments were fully sequenced before any decision was made.

**Figure 1.**
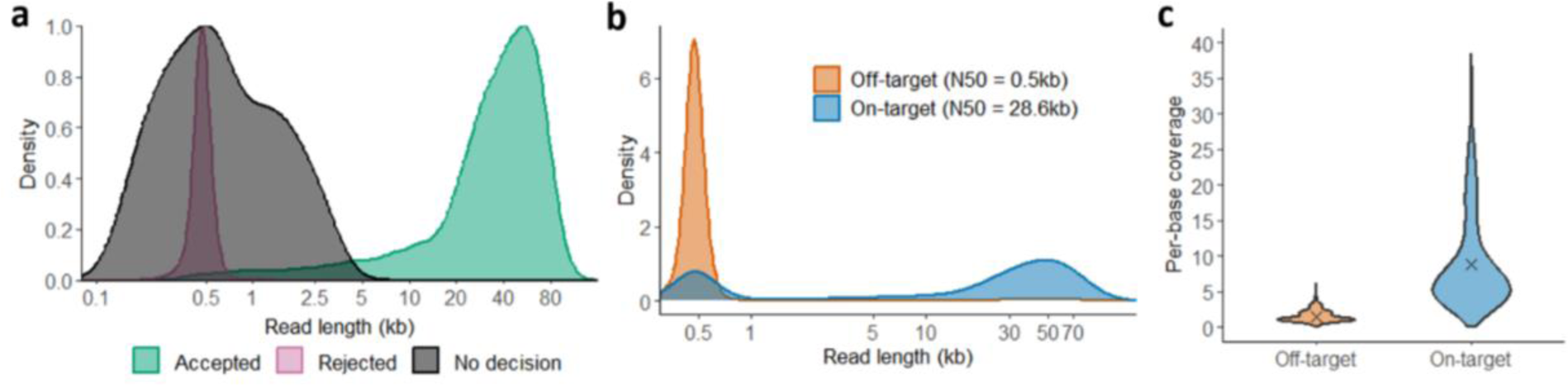
Targeted nanopore sequencing with adaptive sampling on the reference DNA sample HG002. **(a)** Density plot showing read length distribution for different decisions made by the MinKNOW software. The median read lengths of various decisions are: 38,509 nt for accepted reads (green), 471 nt for rejected reads (pink) and 589 nt for reads of no decision (black). **(b)** Density plot showing read length distribution for on-target (blue; N50 = 28.6 kb) versus off-target (orange; N50 = 0.5 kb) alignments. **(c)** Violin plots showing per-base coverage distributions within randomly selected off-target regions (orange) and within on-target regions (blue). A cross (x) marker is incorporated into each violin plot to show the mean coverage of 1.5 and 8.7.

Following the experiment, the Guppy high accuracy (HAC) model was used to re-process the nanopore data and align it to the human hg38 genome. On-target reads exhibited an N50 of 28.6 kb, much higher than the N50 of 500 bps observed for off-target reads (Figure 1b). Furthermore, adaptive sampling resulted in a median 6-fold enrichment in coverage within target regions compared to the off-target regions (Figure 1c). This demonstrated the successful rejection of DNA fragments outside the target regions while fully sequencing DNA fragments within the target regions.

### Adaptive sampling accurately recovers methylation status from control samples

To evaluate the methylation detection with nanopore adaptive sampling, two methylation control samples were sequenced. One with high methylation (>85%), which served as the positive control, and one with low methylation (<5%), which served as the negative control (EpigenDX, 2018). These methylation controls allowed us to evaluate the technology’s accuracy in detecting modified bases from nanopore reads.

The overall methylation levels observed were consistent with the expected methylation level from each control sample (Figure 2a). The mean methylation for the high and low methylation control samples were 93% and 4%, respectively, with approximately 86% of CpG sites fully methylated in the positive control and approximately 94% of sites fully non-methylated in the negative control (Figure 2a). Moreover, for methylation calls only covered by one single read, we observed that the methylation level remained nearly unchanged (Figure 2b). These results are consistent with our previous study indicating that nanopore technology can reliably detect methylation even with just a single read (Yuen et al., 2021). The sequence context of CpG sites detected from our data was then evaluated. For both samples, the four bases around the CpG sites were evenly distributed, indicating no sequence biases in methylation predictions with nanopore sequencing (Figure 2c & d).

**Figure 2.**
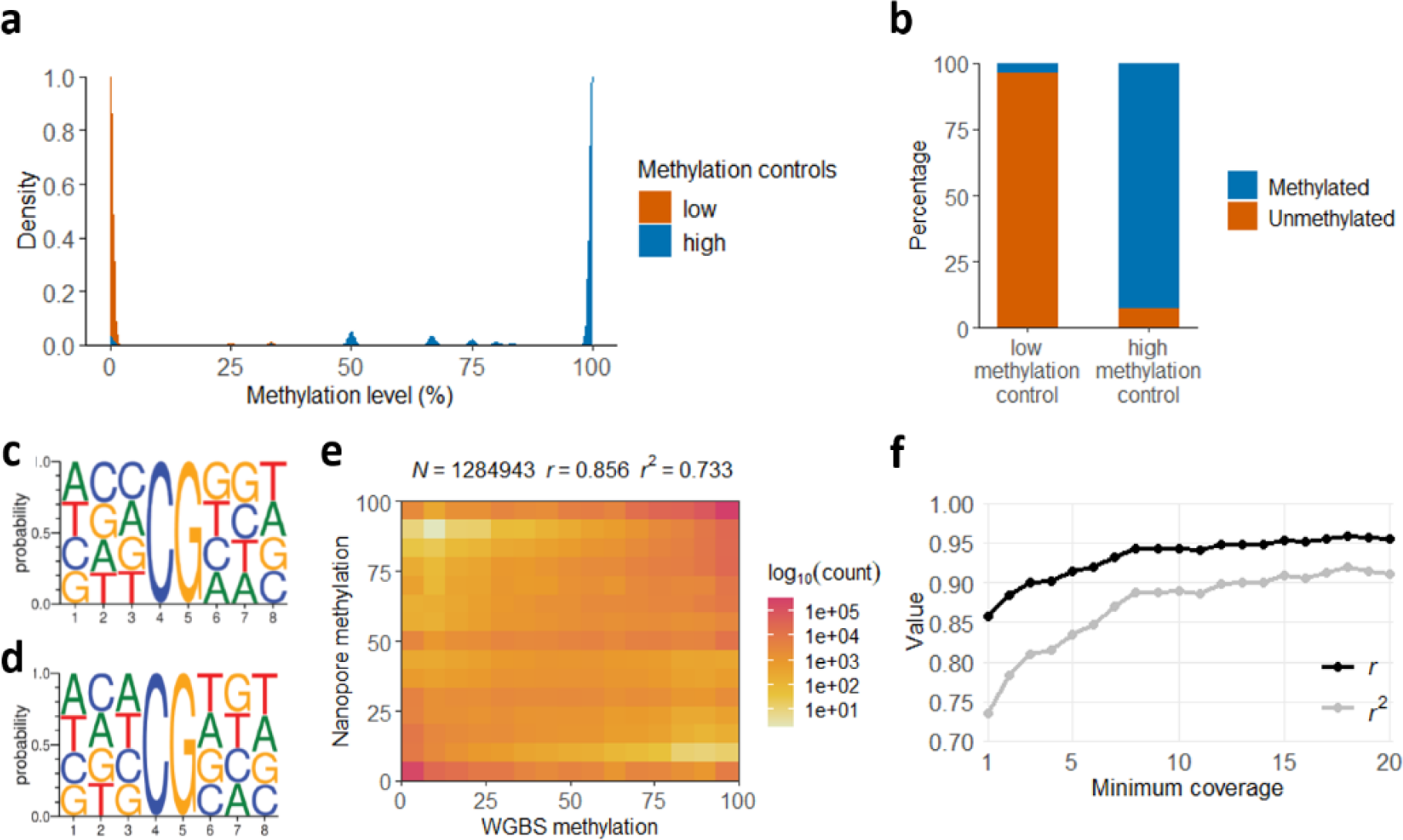
Methylation control samples. **(a)** Density plot showing methylation level distribution for both controls. For the low methylation control, the mean and median methylation are 4% and 0%, respectively (N = 13,593,393). For the high methylation control, the mean and median methylation are 93% and 100% respectively (N = 21,501,907). The expected methylation level reported for these methylation controls by EpigenDX are >85% for high and <5% for low. **(b)** Bar plots showing the proportion of fully methylated (100%) and fully unmethylated (0%) sites, utilising CpG sites with a single read. In the low methylation control, 96% of sites are unmethylated and 4% are methylated, with a total of 7,984,536 sites. In the high methylation control, 93% of sites are methylated and 7% are unmethylated, out of a total of 8,347,590 sites. **(c-d)** Sequence motifs of the DNA 8-mer context of the tested CpG sites (NNNCGNNN) for (c) high methylation control and (d) low methylation control. **(e)** Comparison between nanopore and whole genome bisulfite sequencing (WGBS) for methylation detection. **(f)** Relationship between minimum coverage and correlation metrics (*r* and *r^2^*) for methylation detection using nanopore and WGBS. Each coverage level consists of 1,686 observations. *N* = sample size, *r* = Pearson correlation coefficient, *r^2^* = Coefficient of determination

We also evaluated nanopore methylation against published whole genome bisulfite sequencing (WGBS) data from the HG002 sample. A notable consistency between Nanopore and WGBS data was observed with a Pearson’s correlation coefficient (*r*) of 0.86 (Figure 2e). To determine whether this correlation varied with coverage, we sampled 1,686 observations randomly from each coverage level ranging from 1 to 20 and calculated *r* and *r^2^* values. The correlation values between the WGBS and nanopore estimates showed a slight increase with coverage (Figure 2f). For instance, between coverage >=1 and >=2 reads, the *r*-value increased from 0.86 to 0.89. Considering that the mean coverage of CpG sites with WGBS was 6.9x and with nanopore was 3.3x, we can conclude that nanopore offers the advantage of assessing methylation with fewer reads, while all markers of interest can be analysed simultaneously, and accurately and in a timely manner compared to other MPS methods.

### Nanopore achieves methylation-based body fluid identification

To evaluate methylation markers for body fluid identification, ten human blood samples (Supp. Table 1) were sequenced. To this end, we used a recently developed panel of nine CpG markers to distinguish between semen, blood, vaginal fluid, saliva, and menstrual blood (Lee et al. (2022). For the identification of specific body fluids, the study used the SNaPshot multiplex assay (Lee et al., 2022) and relied on the presence of positive methylation signals at two CpG sites for all sample cases except for saliva, which was based only on a single site, as only one site specific to saliva had been previously identified (Jung et al., 2016)

The nine body fluid specific CpG sites were included in our target panel used in adaptive sampling. By analysing the methylation percentages across the nine sites in each sample, we observed elevated methylation levels specifically at the two target loci for blood (Figure 3), indicating that the methylation measurements obtained from our ten samples using nanopore adaptive sampling accurately assessed these sites as specific markers for blood identification.

**Figure 3.**
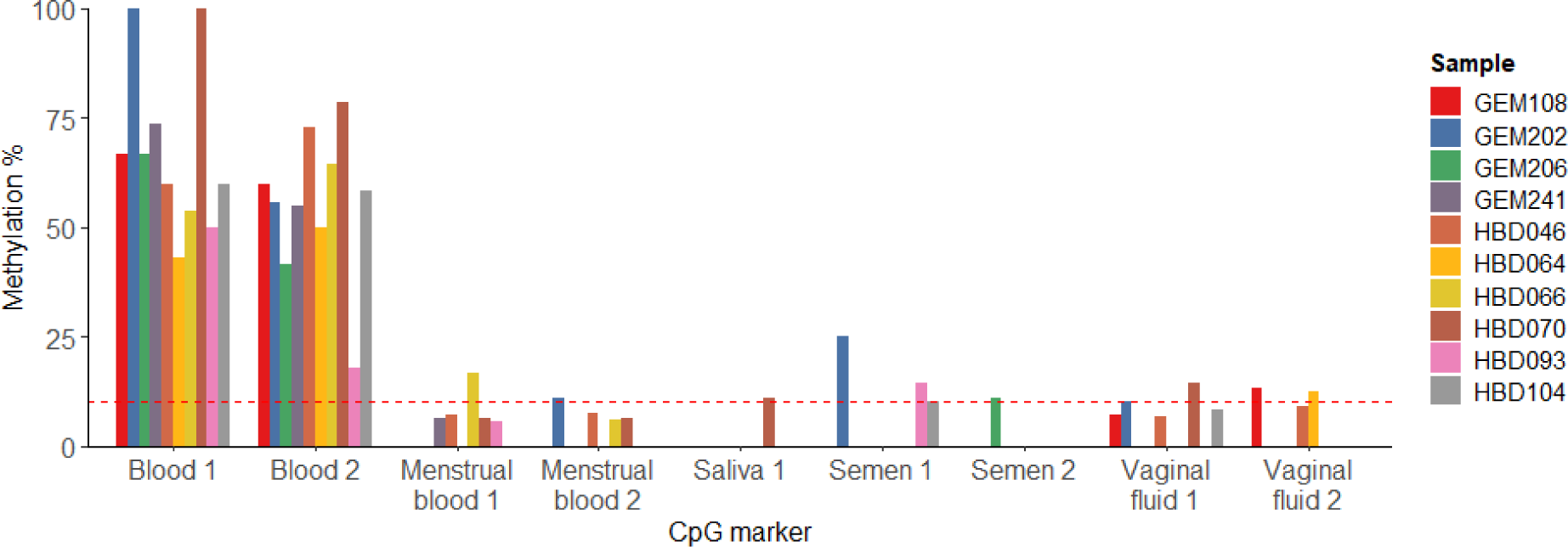
DNA methylation percentage of nine markers used for body fluid identification in ten human blood samples. Each color represents one of the ten human blood samples tested. The y axis indicate the methylation level measured at each of the nine target CpG sites (y axis) for blood (Blood1 – chr19:3344275, Blood2 – chr6:108562706), menstrual blood (Menstrual blood 1 – chr12:57619756, Menstrual blood – chr12:57619734), saliva (Saliva 1 – chr3:194688119), semen (Semen 1 – chr2:219514322, Semen 2 – chr8:143943879), and vaginal fluid (Vaginal fluid – chr7:27251959, Vaginal fluid 2 – chr12:53961752). The dashed red line indicates the 10% threshold suggested by various studies using this panel (Jung et al., 2016; Lee et al., 2022), based on their observed variability of the methylation measurements.

### Assessment of age-associated methylation markers used in forensics

A total of 32 CpGs sites, in eight different genes, that had previously been associated with age were considered (Table 1). These sites are frequently incorporated in the predictive models used to estimate age from DNA samples derived from blood based on multiple forensic studies: Bekaert et al. (2015); Dias et al. (2020); Feng et al. (2018); Fleckhaus and Schneider (2020); Freire-Aradas et al. (2018); Freire-Aradas et al. (2016); Jung et al. (2019); Lee et al. (2022); Naue et al. (2017); Naue et al. (2018); Woźniak et al. (2021); Zbieć-Piekarska, Spólnicka, Kupiec, Makowska, et al. (2015); Zbieć-Piekarska, Spólnicka, Kupiec, Parys-Proszek, et al. (2015). These sites, which we included in our adaptive sampling panel, were used for the assessment of nanopore methylation in our ten samples spanning ages from 25 to 76 (Supp. Table 1). We created a linear regression model commonly used in other forensic methylation studies (Bekaert et al., 2015; Freire-Aradas et al., 2016; Lee et al., 2022; Naue et al., 2018; Woźniak et al., 2021; Zbieć-Piekarska, Spólnicka, Kupiec, Makowska, et al., 2015), to assess the relationship between chronological age from our sample donors and methylation levels at these known CpG sites. A varying degree of correlation was observed, with a majority of CpG sites showing a positive correlation with age and having *r*-value greater than 0.5, indicating that methylation levels at these sites generally increase with age. The site with the highest positive correlation with age was located in the gene *ELOVL2* (at chr6:11044644 (GRCh38)), with an *r*-value of 0.948 (Table 1). While several other sites showed strong positive correlations (*r* ≥ 0.8), a few sites showed weaker correlations with varying values of root mean squared error (RMSE) (Table 1).

**Table 1.**
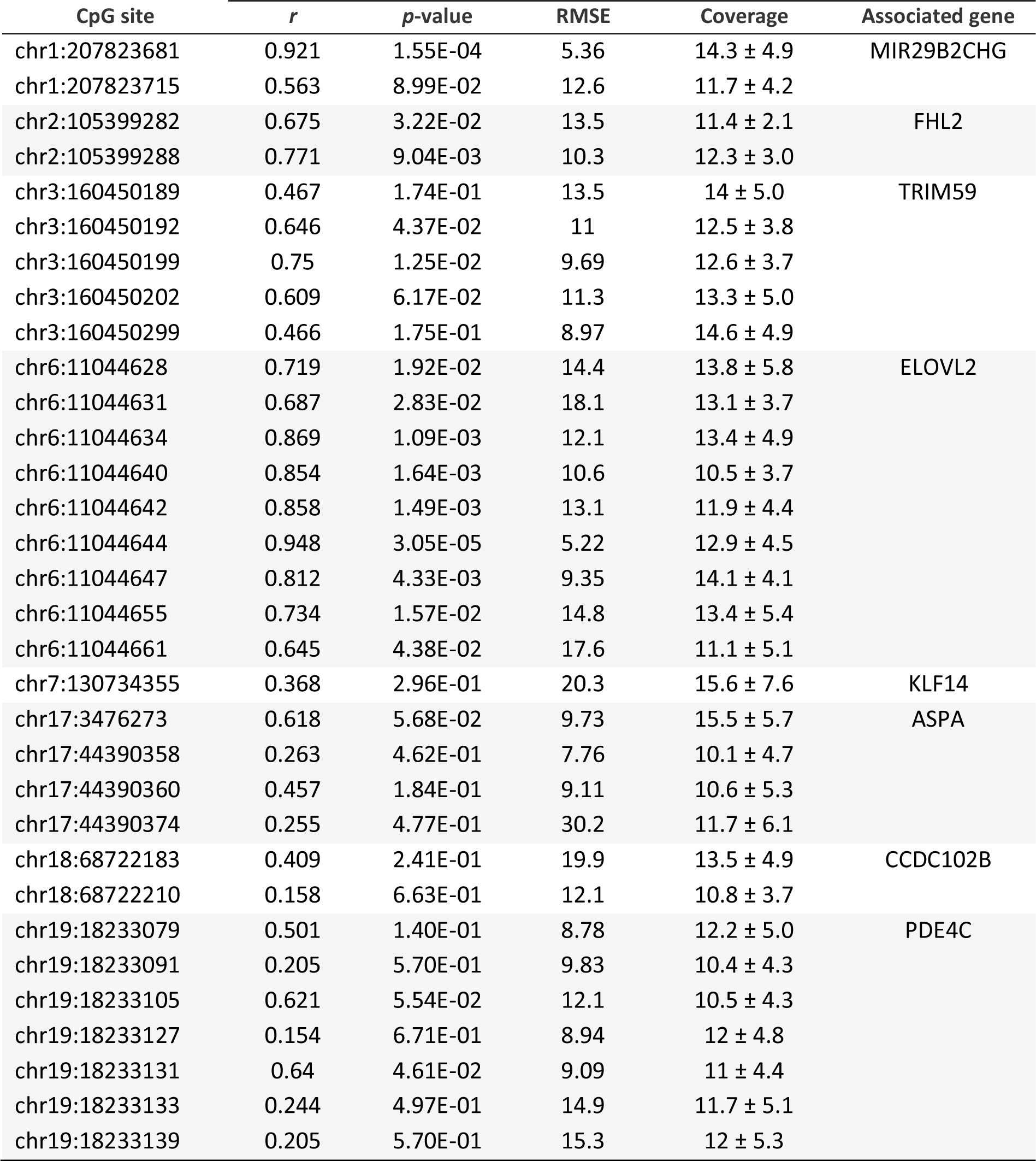
Correlation of chronological age with nanopore-based DNA methylation level for a set of age-prediction markers commonly used in blood-based age prediction models (APMs) in forensics. The CpG sites are located in eight genes (*ELOVL2*, *FHL2*, *MIR29B2CHG*, *CCDC102B*, *KLF14*, *TRIM59,* and *PDE4C*), with the exact chromosomal location (GRCh38/hg38) provided. Simple linear regression was performed between the chronological age of the individuals and methylation levels for each individual CpG site. *r* = Pearson’s R; RMSE = root mean square error; Coverage = mean coverage and standard deviation across ten samples.

### Exploring other age-related genes

Utilizing nanopore adaptive sampling allows for the effortless inclusion of additional target regions in the panel, bypassing the necessity for additional multiplexes, which would normally be required using conventional methodologies. Our targeted design allowed us to explore 307 aging-related human genes from the Aging gene database (Tacutu et al., 2018) alongside 393 orthologous genes that were shown previously to have age-dependent expression across mammals (Wang et al., 2020) (Supp. Table 2). Our study was the first of its kind to use nanopore adaptive sampling to simultaneously detect DNA methylation markers for different forensic purposes in a single assay.

In the aggregated data from the ten tested blood samples, we could measure a total of 2,500,538 CpG sites. We undertook a comprehensive analytical approach to evaluate methylation levels. Following similar analyses previously performed on the forensic age markers (Bekaert et al., 2015; Freire-Aradas et al., 2016; Lee et al., 2022; Naue et al., 2018; Woźniak et al., 2021; Zbieć-Piekarska, Spólnicka, Kupiec, Makowska, et al., 2015), a simple regression analysis was first conducted on this aggregated dataset encompassing all CpG sites identified, yielding a correlation coefficient and associated *p*-value for each site. We then employed the unweighted Stouffer’s method to ascertain the collective statistical evidence across the multiple CpG sites under investigation (Supp. Data 1) (Figure 4a). This method gives a meta Z-score and a meta *p*-value that have the advantage of being comparable across different CpG sites (Lu et al., 2023). Using this approach, we were able to identify numerous CpG sites exhibiting highly significant correlations with age (meta *p*-value < 0.00001) (Figure 4a, highlighted in orange).

**Figure 4.**
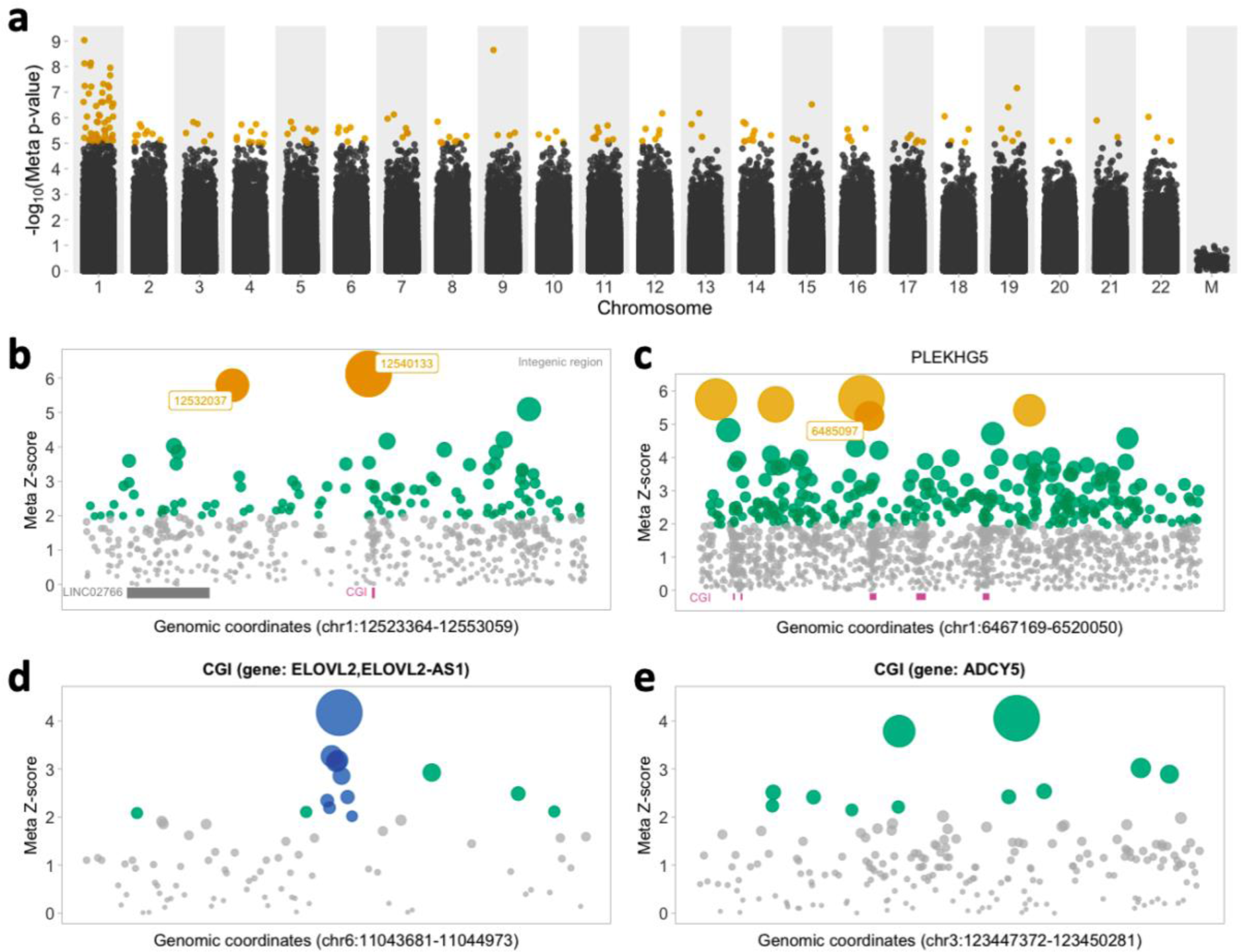
Comprehensive meta-analysis of methylation levels in relation to age. **(a)** The genome-wide distribution of -log_10_(*p*-value) for all CpG sites identified in our sequencing data, plotted across chromosomes except sex chromosomes. CpG sites marked in orange indicate a highly significant correlation with age (meta *p*-value ≤ 0.00001). **(b)** A magnified view of the two highest-significance CpG sites identified in meta-analysis, marked in orange. These CpG sites are chr1:12540133 with a meta *p*-value of 9.26 x 10^-10^ and a meta Z-score of 6.12, followed by the site at chr9:10231889 (data not shown in here) and at chr1:121532037 with a meta *p*-value of 7.08 x 10^-9^. These two sites are in close proximity to each other within intergenic regions. Additional CpG sites potentially linked to age, shown in green, meet the significance threshold with a meta *p*-value of <0.05. The gene *LINC02766*, positioned upstream of the intergenic region, is marked in violet, while a nearby CpG Island (CGI) is marked in blue. CpG sites marked in grey represent those identified in the aggregated sequencing data that do not show a significant association with age. **(c)** A cluster of CpG sites within the *PLEKHG5* gene demonstrate a correlation with age, utilizing a meta *p*-value cutoff of <0.05, indicated in green and orange. CpG sites marked in grey are those detected in the sequencing analysis but do not meet the threshold for significance in relation to age. Specifically, five CpG sites located colored in orange show a strong correlation, each with a meta *p*-value of <1.5 x 10^-7^. Of these, the site chr1:6485097 is positioned within one of the five CGIs identified in the gene. **(d)** Meta-analysis Z-score (Meta Z-score) for CpG sites within the *ELOVL2* gene. Blue points denote CpG sites frequently employed in forensic age prediction models, as detailed in Table 1. Green points denote CpG sites not typically utilized in forensic models but exhibit potential for age prediction applicability, as their meta *p*-value <0.05. **(e)** Analysis identical to that in (d), but on the *ADCY5* gene. Green points show potential for age prediction with meta *p*-value of <0.05. The scatter plots in (b-e) represent the logarithm of the meta *p*-value, where the size of each circle is proportional to the magnitude of the value. Larger circles indicate lower meta *p*-values, denoting higher statistical significance.

Among the top sites that exhibited the most significant association with age, two of them, chr1:12540133 (meta *p*-value = 9.3 x 10^-10^), and chr1:12532037 (meta *p*-value = 7.1 x 10^-9^), were located in an intergenic region near a CpG island and a long non-coding RNA (LINC02766) (Figure 4b, highlighted in orange).

Notably, we also observed five CpG sites within the *PLEKHG5* gene that displayed a high degree of association with age. These sites include chr1:684248 (meta *p*-value = 7.5 x 10^-9^), chr1:6468845 (meta *p*-value = 9.26 x 10^-9^), chr1:6475181 (meta *p*-value = 2.21 x 10^-8^), chr1:6502038 (meta *p*-value = 6.08 x 10^-8^), and chr1:6485097 (meta *p*-value = 1.56 x 10^-7^) (Figure 4c, highlighted in orange). Out of the five CpG islands (CGIs) present in the *PLEKHG5* gene, only one significant CpG site was situated within the CGI. Additionally, a number of other CpG sites in the *PLEKHG5* gene showed an association with aging (meta *p*-value < 0.05) (Figure 4c, highlighted in green). *PLEKHG5* is a protein-coding gene involved in the regulation of neuronal cell differentiation and has not previously been reported to have any known association with age-related changes or diseases.

Nanopore sequencing enables a greater number of CpG sites to be sampled by sequencing entire genes. CGIs, typically situated near transcription start sites, are often associated with regulatory functions of genes and promoter regions. Therefore, the scope was expanded beyond, and around, established age markers, including targeting CpG sites within CGIs, to additional candidates potentially correlating with age. The *ELOVL2* gene, for example, is a well-known predictor of age in blood (Jung et al., 2019; Zbieć-Piekarska, Spólnicka, Kupiec, Makowska, et al., 2015). Nine different CpG sites within this gene have been used and incorporated into different age prediction models (Table 1). Among these nine frequently utilized age-related CpG sites in *ELOVL2*, chr6:11044644 exhibits the strongest association with age in our samples (meta *p*-value = 3.0 x 10^-5^, meta Z-score = 4.2), while chr6:11044661 is the least significant (meta *p*-value = 0.044, meta Z-score = 2.01) (Figure 4d, highlighted in blue). Using a meta p-value of <0.05, we identified six additional sites with potential association with age in *ELOVL2* (meta Z-score > 2.01, meta *p*-value < 0.044) (Figure 4d, highlighted in green).

We subsequently examined other CGIs in our dataset to identify new sites with potentially comparable predictive values. For instance, within the *ADCY5* gene’s CGI, we discovered eleven promising sites that could be useful for predicting age, using the same Z-score threshold applied to the *ELOVL2* gene (meta Z-score > 2.01, meta *p*-value < 0.044) (Figure 4e, highlighted in green).

## Discussion

This study used nanopore adaptive sampling to demonstrate the simultaneous detection of multiple forensic methylation markers for body fluid and age prediction in a single assay, without the need for biochemical treatments of the DNA samples. We corroborated existing markers for body fluids and age using blood samples. By adding multiple target regions in the adaptive sampling sequencing run, we were able to test the genomic regions surrounding the known markers as well as new potential regions of interest, including genic regions plus 20,000nt upstream and downstream. This approach enabled the detection of additional potential methylation markers for age and presented a significant benefit in evaluating methylation using fewer reads, and showing effectiveness even with a single read (Figure 2b & 2f).

When examining age markers commonly used in forensic blood-based models, we found that some sites showed a mild correlation with age (Table 1), suggesting not all markers may be reproducible across different technologies. It is also possible that the number of samples available is not sufficient to recover the age-predictive nature of those sites either through simple regression or using meta-analysis methods, as shown here. Notably, the analysis also lacked replicates entirely. Further evaluation with additional samples for each age range and sample type, i.e. biological source, would assist in improving the statistical power of the tested markers.

The CpG sites chosen for forensic APMs differ from those featured in epigenetic clock models for several reasons. Forensic models primarily aim for precise age prediction of individuals, usually within a specific age variation range. In contrast, epigenetic clocks tend to adopt a more comprehensive approach, emphasizing the biological aging process, which takes into account variations across tissues and maximum lifespan of the species, but with no emphasis on whether predictions are within a specific variation range. Furthermore, the selection of CpG sites is significantly influenced by the types of tissues or cells employed for model testing and validation, the age range of the samples, and the genetic background of the studied population. Not every CpG site linked with chronological age is also associated with biological age, and vice versa.

Despite the variation in the selection of CpG sites between these methods, both models retain their precision and relevance within their designated contexts and applications. As indicated above, the discrepancy in performance might be attributed to the limited number of samples. Nonetheless, the results show that not just a single CpG site from an age-associated gene but rather multiple sites that are close to each other within CGIs could be informative for age prediction. In this regard, alternative sites could be used to add information to the age prediction or to compensate for a site that cannot be used due to the absence of reads. In forensic contexts, this approach is particularly advantageous when working with low template or degraded DNA, offering flexibility and robustness in forensic assay design in terms of selecting specific CpG sites. To consolidate the DNA methylation-age correlation at these CpG sites observed in our study, further tests with larger independent sample sets will be essential.

The integration of the PCR-free long-read approach provided by nanopore sequencing offers benefits in the analysis of forensic evidence, by reducing laboratory preparation steps, reducing PCR artefacts, and increasing the number of markers assessed in a single assay and reducing consumption of the evidence sample through multiple analyses. In particular, it allows genetic and epigenetic markers to be evaluated simultaneously, thereby increasing the informativeness of a single assay. Further, it allows sequencing of native methylated DNA, dispensing with any requirement for a conversion step prior to sequencing and avoiding high DNA input requirements. With nanopore adaptive sampling, targeted sequencing significantly shifts from wet lab manipulations to dry lab computational strategies, making the comprehensive testing of multiple forensic markers more efficient and portable. The recent development of the first epigenetic clock for cattle using nanopore sequencing is a testament to the feasibility of this technology in detecting methylation and developing an APM (Hayes et al., 2021). This advancement highlights that, while nanopore sequencing has been effectively utilized in veterinary science, its potential in constructing similar models for humans remains largely unexplored. This has the potential to transform forensic DNA analysis by making the field-portable nanopore sequencers (e.g., MinION) all-in-one forensic DNA profiling devices for forensic identification and forensic DNA intelligence through simultaneous analysis of biogeographical ancestry, kinship, phenotyping, body-fluid identification and age estimation of the DNA donor.

## Conclusion

This study provides a significant advancement in the field of forensic DNA methylation analysis by employing nanopore adaptive sampling for the simultaneous identification of age-associated and body fluid-specific methylation markers. This innovative approach offers comprehensive profiling of multiple CpG marker sites across entire gene regions from a single sample, eliminating the need for specialized DNA preparation or additional biochemical treatments. The results not only demonstrate that adaptive sampling achieves sufficient coverage and robust consistency with WGBS data, but also corroborates known CpG markers for age and one body fluid (blood). Furthermore, the identification of new sites strongly correlated with age opens up possibilities for discovering novel age methylation markers. Our work paves the way for the systematic development of nanopore-based methodologies, heralding a new era in the field of forensic epigenetic profiling.

## Acknowledgements

We extend our sincere thanks to Dr. Carolina Correa-Ospina and Mr. Lachlan Morrison from the Biomolecular Resource Facility (BRF) at the Australian National University (ANU) for their expert assistance in library preparation and nanopore sequencing.

